# An exploratory study of EEG connectivity during the first year of life in preterm and full-term infants

**DOI:** 10.1101/2021.11.24.469864

**Authors:** Eduardo Gonzalez-Moreira, Deirel Paz-Linares, Lourdes Cubero-Rego, Ariosky Areces-Gonzalez, Thalía Fernández, Manuel Hinojosa-Rodríguez, Pedro A. Valdés-Sosa, Thalia Harmony

## Abstract

**Aim:** To evaluate electroencephalography (EEG) connectivity during the first year of age in healthy full-term infants and preterm infants with prenatal and perinatal risk factors for perinatal brain damage.

**Methods:** Three groups of infants were studied: healthy full-term infants (n = 71), moderate/late-preterm infants (n = 54), and very preterm infants (n = 56). All preterm infants had perinatal and/or perinatal risk factors for brain damage. EEG was obtained during phase II of natural nonrapid eye movement (NREM) sleep. EEG analysis was performed in 24 2.56-s artifact-free segments. For the calculation of EEG sources, spectral structured sparse Bayesian learning was used. Connectivity was computed by the phase-lag index.

**Results:** In healthy full-term infants, EEG interhemispheric connectivity in the different frequency bands followed similar trends with age to those reported in each frequency band: delta connectivity decreased, theta increased at the end of the year, the alpha band showed different trends based on the region studied, and beta interhemispheric connectivity decreased with age. EEG connectivity in preterm infants showed differences from the results of the term group.

**Discussion:** Important structural findings may explain the differences observed in EEG connectivity between the term and preterm groups.

**Conclusion:** The study of EEG connectivity during the first year of age provides essential information on normal and abnormal brain development.

## Introduction

Electroencephalographic brain connectivity in different spectral bands is associated with diverse mechanisms underlying brain development function (1). Band-specific synchronized spectral connectivity is the underlying mechanism for large-scale brain integration of functionally specialized regions from which coherent behavior and cognition emerge (2–5). This mechanism relies mainly on the neural architecture and interactions within layers of cortical columns at the microscopic level of description (6,7). However, in isolation, its topological organization—spatial distribution and connectivity pattern at the macroscopic level possesses tremendous descriptive power with regard to the developing brain of preterm neonates (8).

An essential aspect of describing the development of neural networks involves mapping their spatial distribution alone by the localization of responsive areas at the observational level of experimental techniques, which can be collected by magnetic resonance imaging (MRI) and electroencephalography (EEG) (9). With either approach, this mapping is indirect. However, the spectral composition of functional MRI (fMRI) signals is severely distorted by the slow metabolic-hemodynamic cascade of the process that follows actual neural activity (10,11). Several MRI studies have shown aberrant structural characteristics and even abnormal connectivity in preterm infants (12), suggesting that disrupted white matter tracts may underlie the neurodevelopmental impairments common in this population. It has also been suggested that abnormalities in the functional connectivity between the cortex and thalamus underlie neurocognitive impairments seen after preterm birth (13). The development of thalamocortical connections and how such development relates to cognitive processes during the earliest stages of life, e.g., at ages of one and two years, have been described during the last decade (14).

Furthermore, the thalamus–sensorimotor and thalamus–salience connectivity networks have been shown to be already present in neonates, whereas the thalamus–medial visual and thalamus–default mode network connectivity were shown to emerge later, at one year of age (14). The observation that working memory performance measured at one and two years of age has significant correlations with the thalamus–salience network connectivity is also important. Studies have compared the connectivity between very preterm infants (VPTs) and full-term infants (FTIs) using MRI procedures (15). These studies showed that the most decreased connectivity strength in VPTs was the frontotemporal, fronto-limbic, and posterior cingulate gyrus at gestational ages (GAs) of 39.6 ± 1.2 weeks (FTI) and 40.3 ± 0.6 weeks (VPT).

Although many studies have evaluated structural connectivity by MRI procedures, there is a lack of references using EEG to measure functional connectivity. EEG recordings in neonates and infants have shown that quantitative EEG analyses are a reliable and valuable procedure to evaluate functional and maturational changes (16–18). The study of EEG connectivity is relevant since coherent brain rhythmic activity plays a role in communication between neural populations engaged in functional and cognitive processes (19). It has also been shown that neural synchrony plays a role in synaptic plasticity (20). Therefore, the study of EEG connectivity in preterm and term infants early in life may provide essential knowledge regarding brain development. However, EEG signals are affected by their low spatial resolution and volume conduction effects (21,22). These pitfalls have been addressed by deploying generations of electrophysiological source imaging (ESI) metho2ds during recent decades (23). ESI methods combine the best spatial resolution of MRI for head model estimation with the better time resolution of EEG for inference of neural activity and connectivity at the brain level (24–28).

The World Health Organization estimates the prevalence of preterm birth to be 5–18% across 184 countries worldwide (29). The causes for premature birth comprise mainly biological, genetic, and environmental factors (30). Despite advances in prenatal and neonatal care and decreased perinatal mortality of preterm newborns, the number of survivors with neurological and cognitive deficits constitutes a public health problem (31). Furthermore, preterm birth is a leading risk factor for (i) cerebral palsy (32,33), (ii) delayed mental and/or psychomotor development (34,35), (iii) executive dysfunction (36), (iv) neurosensory disability (37), (v) language and reading deficits (38), (vi) academic underachievement (39,40), (vii) attention-deficit/hyperactivity disorder, and (viii) autism spectrum disorders (12,41).

In this study, we focused on the temporal dynamics of neural networks in the millisecond range to study early neural integration. We present a longitudinal study of EEG connectivity in preterm infants and FTIs during the first year of life using a measure based on instantaneous phase differences.

## 1 Methods

Ethical permission was granted by the Ethics Committee of the Instituto de Neurobiología of the Universidad Nacional Autónoma de México, which complies with the Ethical Principles for Medical Research Involving Human Subjects by the Helsinki Declaration. Informed consent from the parents was obtained for all study participants.

### 1.1 Participants

Three groups of infants were studied: i) healthy FTIs without any antecedent for perinatal brain damage; ii) a group of moderate/late-preterm infants with GA between 32 and 37 weeks; and iii) a group of VPTs with a GA of 27 to 31 weeks. All preterm babies had prenatal and/or perinatal risk factors for perinatal brain damage. However, participants with congenital and hereditary brain malformations or infectious or parasitic diseases were excluded from this study. After the infants were discharged from the hospital where they were born, their parents were invited to participate in a unique project of the Neurodevelopmental Research Unit at the Institute of Neurobiology of the National Autonomous University of Mexico in Queretaro. Information regarding each group is included in Table 1.

**Table 1.** Demographic information for full-term and preterm infants.

| Age Group | N | Sex (females) | EEGs | GA (wks) |
| --- | --- | --- | --- | --- |
| Full-term | 71 | 29 | 82 | 40 (38-41) |
| Moderate/late preterm | 54 | 25 | 112 | 35 (32-37) |
| Very preterm | 56 | 27 | 103 | 29 (27-31) |

### 1.2 EEG data analysis

EEG was acquired from infants while they were in phase II sleep and were in his or her mother’s lap in a dimly lit room with acoustic isolation. No sedation was used. Referential EEG recordings for 20 minutes were obtained from 19 electrodes based on the 10/20 system using linked ears as a reference. The MEDICID IV System with a gain of 20,000, amplifier bandwidth between 0.5 and 100 Hz, and sample rate of 200 Hz was used. Some participants were recorded two or more times during their first year of life. Therefore, EEG data were selected from a dataset of 297 recordings collected between 2016 and 2020 as part of an ongoing project investigating the characterization of brain development in preterm infants.

Later, EEG data were segmented by visual inspection into 24 artifact-free segments of 2.56-s duration. The idea behind this processing was to avoid transforming original data by using preprocessing techniques such as independent component analysis to remove artifacts (eye movement and blinks) or interpolation to fix “bad” channels. These methods could profoundly modify the connectivity information relayed in the electrophysiology signal, leading us to an incorrect interpretation of the data. The frequency analysis was set up from 0.3 Hz to 20 Hz. After EEG data collection and selection, the data analysis continued with two main steps: inference of EEG source space data using an ESI method and connectivity analysis based on the phase-lag connectivity measure.

### 1.3 Inference of EEG source space data

ESI methods aim to infer local neural currents based on EEG and MRI data (42,43). However, this process is subject to distortions due to the system of linear equations being highly ill-conditioned and because the possible solutions lie within a high-dimensional space. These distortions can reach unacceptable levels as repeatedly shown in simulations (21,22,44).

In this paper, we estimated cortical neural activity using a third generation ESI method, spectral structured sparse Bayesian learning (sSSBL) (45). sSSBL pursues estimation of neural activity through a maximum “evidence” search via the expectation-maximization algorithm (46). Evidence is defined as the conditional probabilities of two groups of parameters: (i) variances in spectral EEG source activity, which controls the statistical relevance of the source cross-spectral components; and (ii) variances in spectral EEG noise, which controls the level of noise in the observations.

Furthermore, this approach is based on an iterative scheme that produces an approximated representation of the evidence (expectation) followed by its maximization, guaranteeing convergence to a local maximum. The maximization step is carried out via estimation formulas of the vector regression elastic net (47) and sparse Bayesian learning (48) through the arithmetic mean of typical vector regression inputs corresponding to the samples. The global sparsity level is handled by estimating the regularization parameters in a completely analogous form to the procedure described by Paz-Linares and collaborators (47).

The cortical activity inference was set up on a cortical manifold space defined as 5000 points at the gray matter, with coordinates on the pediatric Montreal Neurological Institute (MNI) brain template (http://www.bic.mni.mcgill.ca). The scalp sensor space was built on 19 electrodes within a 10-20 EEG sensor system (49). The lead fields were computed by the boundary element method (BEM) integration method accounting for a model with five head compartments (gray matter, cerebrospinal fluid, inner skull, outer skull, scalp) (50). The initial cortical surface parcellation based on ninety regions of Tzourio-Mazoyer’s atlas (51) was manually categorized into five large regions per hemisphere: frontal, sensorimotor, parietal, temporal, and occipital regions.

### 1.4 Connectivity analysis through phase-lag based measures

In neuroscience, phase locking has become the primary measure for neural connectivity to evaluate the synchronization between neural groups. In this study, we computed the phase-lag index (PLI) (52,53) between regions of interest (ROIs). The PLI is one of the most popular methods for synchronization inference because of its near “immunity” against volume conduction effects (54). This approach applies spatial filters to EEG data that reduce volume conduction effects, leading to the correct interpretation of connectivity information.

In this study, an all-to-all PLI connectivity matrix was computed between the ten cortical regions, five per hemisphere, at each frequency point from 0.3 Hz to 20 Hz. This procedure resulted in a 3-D matrix (ROI-ROI-frequency). Finally, global efficiency was computed to assess the connectivity information per cortical region and to summarize the connectivity matrices. Efficiency is based on the inverse of the average distance from each vertex (ROI) to any other vertex (path lengths), which explains why higher efficiency values correspond to more direct connections. Furthermore, global efficiency has been proven to be helpful in evaluating pathol1.5 ogical networks since it is robust against networks that are not fully connected (55,56).

### 1.5 Statistical analysis

For the statistical analysis, a 3-D efficiency matrix (ROI-frequency-age) was created for each group under analysis. One point to note is that our data did not cover every age value between 0 and 1 year. Figure 1 shows some small gaps in the age distribution of each group. To address this pitfall and to estimate with more critical details the development connectivity surfaces during the first year of life, a locally weighted scatterplot smoothing (LOWESS) method was applied (57). The LOWESS approach overcomes classical methods through a linear and nonlinear least squares regression. This regression fits simple models with subsets of the data to build up a function that describes the deterministic part of the variation in the data instead of requiring a global function to fit a model to the entire dataset. Later, we computed a linear regression model for each row of the development connectivity surface to evaluate the connectivity behavior for each frequency value during the first year of life. The slope-based curve provides evidence to compare the connectivity developed for the three groups under analysis.

**Fig 1:**
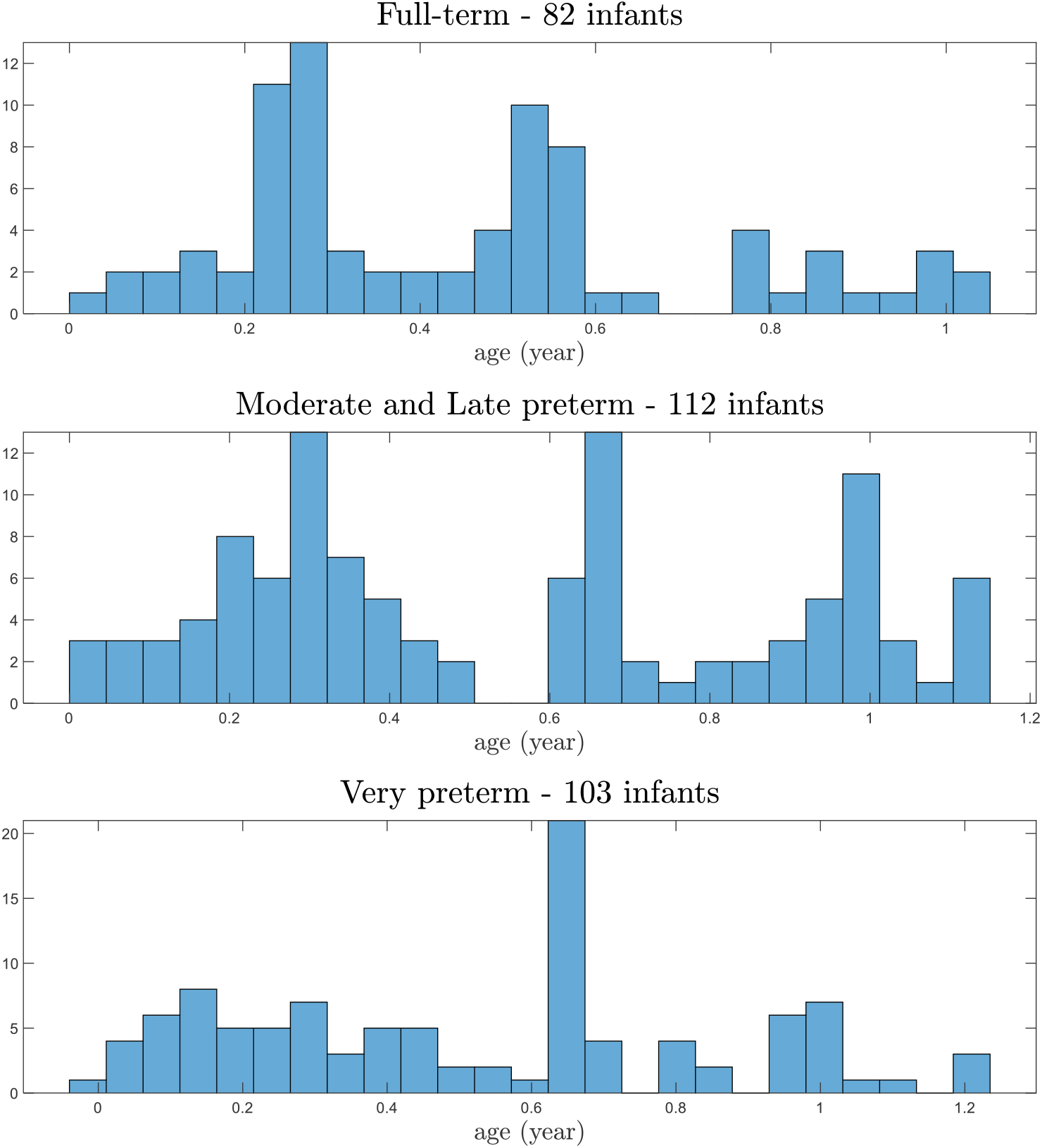
Histogram by age groups showing some small gaps in EEG data recordings during the first year.

## 2 Results

The results of the LOWESS approach for global efficiency measurements in FTIs and preterm infants are shown in Figure 2a. In this figure, the term group showed a decrease in connectivity with age within the delta band frequencies (0.5 Hz–3.0 Hz), whereas in both groups of preterm infants, an increase was observed. Connectivity in the theta and alpha bands decreased in the three groups. In the low beta band, EEG connectivity increased in the term and very preterm groups and decreased in the moderate/late-preterm group. Connectivity at frequencies at 15 Hz and above decreased in all groups. As it is difficult to interpret the results of global connectivity power and almost all previous studies studied interhemispheric connectivity, we present the results based on interhemispheric connectivity for each cortical region under analysis.

**Fig 2:**
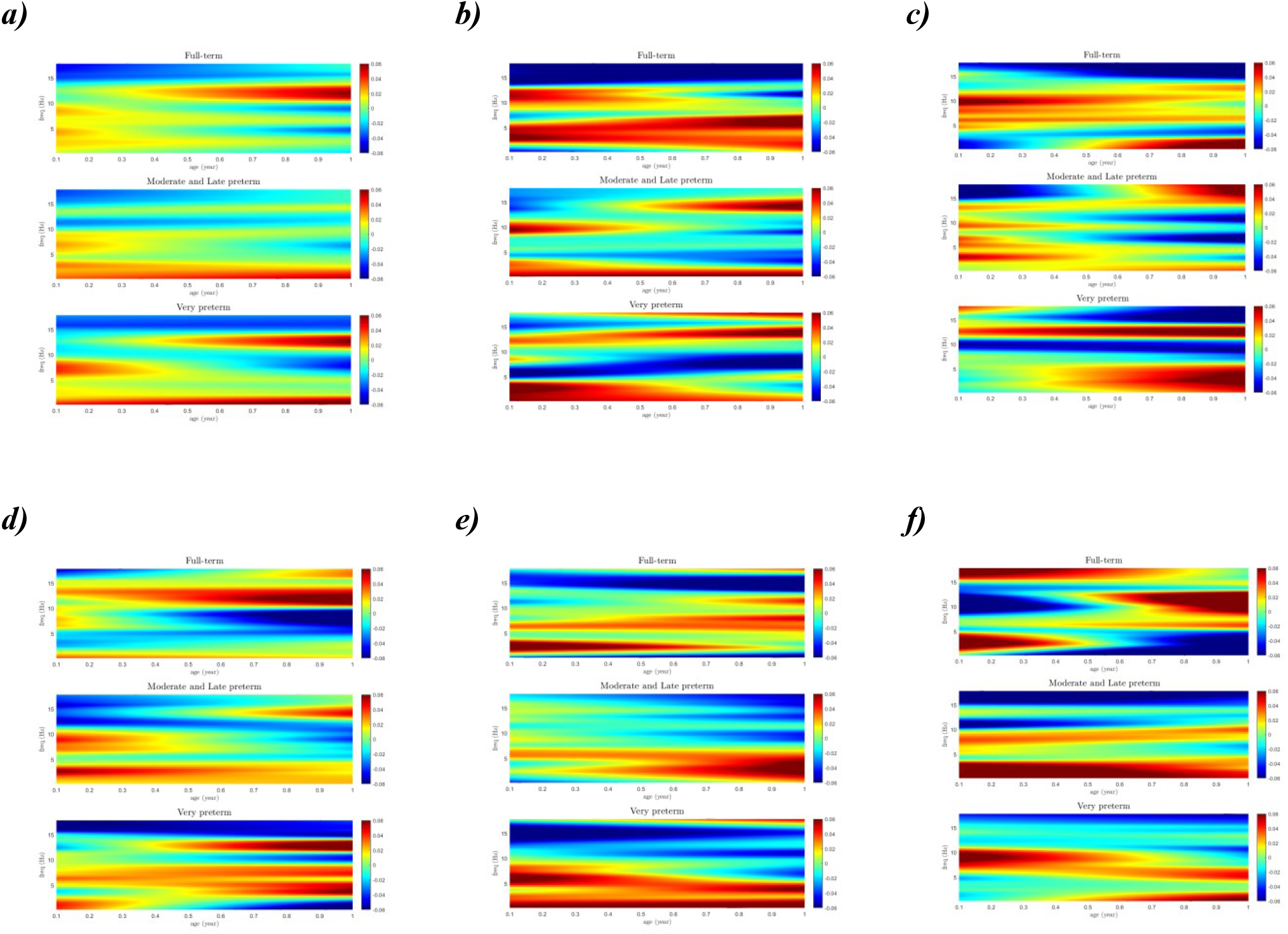
Connectivity patterns in the three groups of infants: full-term, moderate/late preterm, and very preterm: a) global connectivity patterns, b) interhemispheric frontal cortical connectivity patterns, c) interhemispheric sensorimotor cortical regions connectivity patterns, d) interhemispheric parietal cortical regions connectivity patterns, e) interhemispheric temporal cortical regions connectivity patterns, f) interhemispheric occipital regions connectivity patterns.

### 2.1 Left-right frontal connectivity (LRFC)

LRFC results are shown in Figure 2b. In the delta band, LRFC tended to decrease, although at age 0.1 years (36.5 days), different connectivity values were observed in different delta band frequencies. In the moderate/late-preterm group, this interhemispheric delta connectivity decreased, although at 0.5 Hz–1.0 Hz there was an increase with age. In the very preterm group, a decrease in interhemispheric frontal connectivity with increasing age was evident. LRFC in the theta band showed an increase in full-term subjects with age. Meanwhile, in the moderate/late-preterm group, there was a decrease in connectivity with age that was more marked in the very preterm group. LRFC in the alpha band in term and moderate/late-preterm infants decreased with age. As power in this frequency band decreased, this result was expected. However, there were differences in connectivity trends between groups in the beta band: full-term infants had a decreasing trend during the first year of life, but both groups of preterm infants showed increasing values with age.

### 2.2 Left-right sensorimotor connectivity (LRSC)

These results are shown in Figure 2c. LRSC in the delta band in term infants decreased during the first six months of age and subsequently increased. At the end of the first year, this increase was also observed in the moderate/late-preterm infants and with great intensity in the VPTs. This latter group also showed this increase in LRSC at frequencies in the theta band; meanwhile, a decrease in LRSC in the theta band was observed in full-term infants and in moderate/late-preterm infants. There was a constant increase in the 5 Hz–8 Hz range in term infants. This may correspond to rhythmic central activity that has been reported as a precursor to the mu or sensorimotor rhythm (58). Moderate/late-preterm infants showed a decrease in theta connectivity, and theta connectivity in VPTs showed a constant decrease. In the alpha band, LRSC decreased with age in term and moderate/late-preterm infants. In both groups at three months, robust alpha connectivity slowly decreased. However, in the VPTs, there was a substantial decrease across the whole year. LRSC at approximately 15 Hz decreased across the whole year in term infants and VPTs, and there was an unexpected increase at the end of the year in the moderate/late-preterm infants.

### 2.3 Left-right parietal connectivity (LRPC)

The results regarding LRPC are shown in Figure 2d. LRPC in term infants showed a decrease with age at all frequencies between 5 Hz and 10 Hz. However, in moderate/late-preterm infants, LRPC in the delta frequencies and the band from 6 Hz to 10 Hz showed high values at 3.65 months of age that progressively decreased with increasing age. In this group, connectivity in the band from 11 Hz to 18 Hz decreased, but connectivity had increased with age at the end of the year.

### 2.4 Left-right temporal connectivity (LRTC)

Figure 2e shows the results for LRTC. In the delta band, this connectivity at three months was robust and decreased with age in term infants. Moderate/late-preterm infants and VPTs showed the reverse trend, i.e., increases with age. In term infants, LRTC at frequencies from 6 to 8 Hz increased during the year. At 10 Hz to 12 Hz, LRTC decreased with age up to 6 months and subsequently increased. LRTC at frequencies within the beta band decreased with age in term infants. LRTC in both preterm groups decreased with age in the band from 10 Hz to 18 Hz.

### 2.5 Left-right occipital connectivity (LROC)

In Figure 2f, the results regarding LROC are shown. FTIs showed a progressive decrease with age in delta band frequencies. In the alpha band, connectivity decreased during the first six months and increased in the last six months of the year. Moderate/late-preterm infants and VPTs showed completely different patterns of connectivity behavior across the first year of age.

## 3 Discussion

In this study, EEG connectivity in preterm infants was described. Most papers reporting brain connectivity in preterm infants have used MRI (30,59–61), and prominent differences between networks identified in term control versus premature infants at term-equivalent ages have been described (62). These authors also reported that putative precursors of the default mode network were detected in term control infants but were not identified in preterm infants, including those at term-equivalent ages. In a follow-up study of preterm children at seven years of age, (63) demonstrated that children born very preterm had less connected and less complex brain networks than typically developing term-born children and that these structural abnormalities were observed after a follow-6up of seven years. Structural information about the connectivity observed in preterm infants demonstrated alterations related to motor, linguistic and cognitive deficits (64). All this information was the basis for studying EEG connectivity in preterm infants.

The pioneering studies of (65) reported EEG coherence from eight left and eight right intrahemispheric electrode pairs from 253 children ranging in mean age from 6 months to 7 years. The results supported the view that the functions of the left and right hemispheres are established early in human development through complementary developmental sequences. These sequences appeared to recapitulate differences in adult hemispheric function. However, subsequent studies in infants have mainly analyzed the correlation between homologous left and right hemispheres.

Previous studies analyzing the correlations between homologous left and right hemispheres (66) have described that median correlation values significantly decreased (between -40% and -60% decrease) in infants from 27 to 37 weeks of GA. Regarding postnatal maturation, only the central-temporal channel showed a decreasing trend. These authors concluded that the decreasing median correlation values in all homologous channels indicated a decrease in similarity in signal shape with increasing GA. González et al., in 2011, studied EEG inter- and intrahemispheric connectivity measuring coherence between regions and a measure of phase synchronization (67). They found significant differences between term and preterm infants during active and quiet sleep, with term infants having greater magnitude values of coherence than preterm infants. The interhemispheric PLI values were different during active sleep between term and preterm infants in the delta band. Similarly, the intrahemispheric PLI values in the beta band differed between term and preterm infants during quiet sleep. Our results showed that full-term infants had different EEG connectivity results during quiet sleep than preterm infants in all frequency bands. Differences in the EEG analysis may explain the contradiction with the González results.

Significant structural differences may explain the findings observed in EEG connectivity between the term and preterm groups. Several studies have shown that preterm infants with risk factors for perinatal brain damage, especially those born before 32 gestational weeks, exhibit cerebral white and gray matter abnormalities (31). For example, brain MRI studies report that 50 to 80% of extreme and very premature infants present mild-to-moderate abnormalities in the cerebral white matter (68). Myelination delays in the posterior limb of the internal capsule and/or corona radiata, enlargement of the lateral ventricles (due to cerebral white matter loss) and thinning of the corpus callosum (CC) have frequently been observed on MRI. This last neuroradiological finding can range from a partial thinning of the CC with a decrease in the isthmus and/or body to a global thinning in the CC (69). It is important to mention that the CC is a cerebral structure constituted by axons connecting homologous cortical regions. Rapid growth in its volume occurs during the first 20 months of age (68,70). The midsagittal area of the CC has been commonly used as a sensitive marker of brain development and maturation since the area of the CC is related to the number of axons and their morphology, such as axon diameter and myelination (71). Furthermore, the loss of CC at the level of the isthmus and the mid-body has been associated with alterations in sensorimotor interhemispheric connectivity due to the loss of transcallosal connections, which affects maturation of the motor control network (72,73). This spectrum of structural abnormalities may explain many differences noted between term and preterm infants in interhemispheric EEG connectivity, especially in the sensorimotor region. Therefore, the development of the corpus callosum in preterm infants is affected by prematurity (74), and in preterm infants, a decrease in its volume has frequently been observed (69). This structural abnormality may explain many differences noted between term and preterm infants in the interhemispheric EEG connectivity, which we consider the leading cause of the resu7lts obtained.

Another important aspect is that cortical synaptogenesis shows different patterns of development across the cortex, with a more rapid increase in the auditory cortex than in the prefrontal cortex (75), which may explain the asynchrony in cortical maturation in the infant’s brain (76). These facts, together with the maturational process of myelination, show that it ends at a different time in different regions: auditory and visual cortex myelination ends at 18-24 months, whereas these processes end in Broca’s area at five years and in the prefrontal cortex at nine years of age (77). These findings may explain essential differences in the topography of EEG connectivity during the first year of age. On the other hand, myelination in preterm babies is severely affected since MRI studies have shown that diffuse white matter injury is one of the most frequent abnormalities observed in preterm infants (78). The structural differences between term and preterm babies strongly support the differences observed between this group in EEG connectivity.

In the group of term infants, the results obtained may be explained by studies of EEG development in normal infants (79). In all regions studied, EEG connectivity in the delta band decreased with age. EEG development in this frequency band has also been shown to decrease with age, which may explain the results observed in connectivity. EEG connectivity in the theta band shows differences in development based on the region being studied. LRFC showed a significant increase with age, which is consistent with the observation that in term infants’ theta, absolute and relative power in frontal leads increase during the first year (79). LRTC also showed an increase at the end of the year, which coincided with the EEG neurodevelopmental findings.

In the range from 5 Hz to 8 Hz in FTIs, there was a constant increase in LRFC, as well as in LRTC and LRSC. This may correspond to rhythmic central activity that has been reported as a precursor to the mu or sensorimotor rhythm (58). The moderate/late-preterm group showed a decrease in connectivity, and in the very preterm group, this activity constantly decreased. Our findings are consistent with (80). A clear sensorimotor rhythm was described in the range of 5.47–7.03 Hz with contralateral activity in response to free movement while awake FTIs around the four months of life. Furthermore, the preterm infant group with periventricular leukomalacia did not show any electroencephalographic sign of the presence of this rhythm.

In the alpha band, EEG connectivity in term infants showed different trends in the different regions. In the frontal, temporal, and sensorimotor regions, interhemispheric connectivity decreased with age across the whole year. However, LROC showed a sharp decrease in the early months that changed to a progressive increase in the second semester of the year. This changing trend was not detected in the studies of EEG development, perhaps because they used linear regression for the analysis (79).

EEG beta band connectivity decreased in all regions. Our results were limited to a small range of frequencies, from 13 to 20 Hz. Therefore, it is difficult to compare with other studies of EEG development.

## 4 Conclusions

Our exploratory study of EEG connectivity between left and right cortical areas in healthy FTIs during the first year of life showed a similar trend that has been reported in the different frequency bands in similar groups of healthy FTIs. EEG interhemispheric connectivity in all preterm infants studied with a GA from 26 to 37 weeks and prenatal and perinatal risk factors for brain damage showed great differences from the group of healthy FTIs. Such differences in EEG connectivity may be due to the structural brain abnormalities that have been described in preterm infants.

## 6 Author Contributions

TH conceived the project and contributed to the design of the study and validations. EGM contributed to the design of the study, data analysis, validations, and drafted the figures. DPL and AAG contributed to the design of the software tool implementation, carried out computational experiments and validations. LCR, MHR and PAVS contributed to the design of the study and supervised validations. All authors contributed to the final manuscript.

## 8 Acknowledgments

This work received support from Luis Aguilar, Alejandro de León, and Jair García from Laboratorio Nacional de Visualización Científica Avanzada. The authors would also like to thank colleagues Hector Belmont, María Elizabeth Monica Carlier, and María Elena Juarez for their support during data collection. We also acknowledge the collaboration of the Escuela Nacional de Trabajo Social and the participation of Marcela García-Tinoco.

## References

1. Brookes MJ, O’Neill GC, Hall EL, Woolrich MW, Baker A, Palazzo Corner S, et al. Measuring temporal, spectral and spatial changes in electrophysiological brain network connectivity. NeuroImage [Internet]. 2014 May;91:282–99. Available from: https://linkinghub.elsevier.com/retrieve/pii/S1053811914000123

2. Engel AK, Fries P, Singer W. Dynamic predictions: Oscillations and synchrony in top– down processing. Nature Reviews Neuroscience [Internet]. 2001 Oct;2(10):704–16. Available from: http://www.nature.com/articles/35094565

3. Varela F, Lachaux J-P, Rodriguez E, Martinerie J. The brainweb: Phase synchronization and large-scale integration. Nature Reviews Neuroscience [Internet]. 2001 Apr [cited 2021 Oct 11];2(4):229–39. Available from: http://www.nature.com/articles/35067550

4. Nunez PL, Srinivasan R. Electric Fields of the Brain [Internet]. Oxford University Press; 2006. Available from: https://oxford.universitypressscholarship.com/view/10.1093/acprof:oso/9780195050387.001.0001/acprof-9780195050387

5. Vidaurre D, Abeysuriya R, Becker R, Quinn AJ, Alfaro-Almagro F, Smith SM, et al. Discovering dynamic brain networks from big data in rest and task. NeuroImage [Internet]. 2018 Oct 15;180:646–56. Available from: https://linkinghub.elsevier.com/retrieve/pii/S10538119173054879

6. Freeman WJ. Mass action in the nervous system [Internet]. 1975. Available from: www.ccs.fau.edu/~bressler/EDU/NTSA/References/MASS

7. Jirsa VK, Haken H. A derivation of a macroscopic field theory of the brain from the quasi-microscopic neural dynamics. Physica D: Nonlinear Phenomena [Internet]. 1997 Jan;99(4):503–26. Available from: https://linkinghub.elsevier.com/retrieve/pii/S0167278996001662

8. Wilson-Costello D, Friedman H, Minich N, Fanaroff AA, Hack M. Improved survival rates with increased neurodevelopmental disability for extremely low birth weight infants in the 1990s. Pediatrics. 2005 Apr;115(4):997–1003.

9. Mantini D, Perrucci MG, del Gratta C, Romani GL, Corbetta M. Electrophysiological signatures of resting state networks in the human brain. Proceedings of the National Academy of Sciences [Internet]. 2007 Aug 7;104(32):13170–5. Available from: http://www.pnas.org/cgi/doi/10.1073/pnas.0700668104

10. Buxton RB, Wong EC, Frank LR. Dynamics of blood flow and oxygenation changes during brain activation: The balloon model. Magnetic Resonance in Medicine [Internet]. 1998 Jun;39(6):855–64. Available from: https://onlinelibrary.wiley.com/doi/10.1002/mrm.1910390602

11. Logothetis NK, Pauls J, Augath M, Trinath T, Oeltermann A. Neurophysiological investigation of the basis of the fMRI signal. Nature [Internet]. 2001 Jul;412(6843):150–7. Available from: http://www.nature.com/articles/35084005

12. Rogers CE, Lean RE, Wheelock MD, Smyser CD. Aberrant structural and functional connectivity and neurodevelopmental impairment in preterm children. Journal of Neurodevelopmental Disorders [Internet]. 2018 Dec 13;10(1):38. Available from: https://jneurodevdisorders.biomedcentral.com/articles/10.1186/s11689-018-9253-x

13. Toulmin H, O’Muircheartaigh J, Counsell SJ, Falconer S, Chew A, Beckmann CF, et al. Functional thalamocortical connectivity at term equivalent age and outcome at 2 years in infants born preterm. Cortex [Internet]. 2021 Feb 1;135:17–29. Available from: https://linkinghub.elsevier.com/retrieve/pii/S001094522030366X

14. Alcauter S, Lin W, Smith JK, Short SJ, Goldman BD, Reznick JS, et al. Development of Thalamocortical Connectivity during Infancy and Its Cognitive Correlations. Journal of Neuroscience [Internet]. 2014 Jul 2;34(27):9067–75. Available from: https://www.jneurosci.org/lookup/doi/10.1523/JNEUROSCI.0796-14.2014

15. Sa de Almeida J, Meskaldji D-E, Loukas S, Lordier L, Gui L, Lazeyras F, et al. Preterm birth leads to impaired rich-club organization and fronto-paralimbic/limbic structural connectivity in newborns. NeuroImage [Internet]. 2021 Jan 15;225:117440. Available from: https://linkinghub.elsevier.com/retrieve/pii/S1053811920309253

16. Niemarkt HJ, Andriessen P, Peters CHL, Pasman JW, Zimmermann LJ, Bambang Oetomo S. Quantitative analysis of maturational changes in EEG background activity10in very preterm infants with a normal neurodevelopment at 1year of age. Early Human Development [Internet]. 2010 Apr;86(4):219–24. Available from: https://linkinghub.elsevier.com/retrieve/pii/S0378378210000629

17. Niemarkt HJ, Jennekens W, Pasman JW, Katgert T, van Pul C, Gavilanes AWD, et al. Maturational Changes in Automated EEG Spectral Power Analysis in Preterm Infants. Pediatric Research [Internet]. 2011 Nov;70(5):529–34. Available from: http://www.nature.com/doifinder/10.1203/PDR.0b013e31822d748b

18. Schaworonkow N, Voytek B. Longitudinal changes in aperiodic and periodic activity in electrophysiological recordings in the first seven months of life. Developmental Cognitive Neuroscience [Internet]. 2021 Feb 1;47:100895. Available from: https://linkinghub.elsevier.com/retrieve/pii/S1878929320301420

19. Wang X-J. Neurophysiological and Computational Principles of Cortical Rhythms in Cognition. Physiological Reviews [Internet]. 2010 Jul;90(3):1195–268. Available from: https://www.physiology.org/doi/10.1152/physrev.00035.2008

20. Uhlhaas PJ, Roux F, Rodriguez E, Rotarska-Jagiela A, Singer W. Neural synchrony and the development of cortical networks. Vol. 14, Trends in Cognitive Sciences. 2010. p. 72–80.

21. Haufe S, Nikulin V v., Müller K-R, Nolte G. A critical assessment of connectivity measures for EEG data: A simulation study. NeuroImage [Internet]. 2013 Jan 1;64(1):120–33. Available from: https://linkinghub.elsevier.com/retrieve/pii/S1053811912009469

22. van de Steen F, Faes L, Karahan E, Songsiri J, Valdes-Sosa PA, Marinazzo D. Critical Comments on EEG Sensor Space Dynamical Connectivity Analysis. Brain Topography [Internet]. 2019 Jul 30;32(4):643–54. Available from: http://link.springer.com/10.1007/s10548-016-0538-7

23. Gonzalez-Moreira E, Paz-Linares D, Areces-Gonzalez A, Wang R, Valdes-Sosa PA. Third Generation MEEG Source Connectivity Analysis Toolbox (BC-VARETA 1.0) and Validation Benchmark. 2018 Oct 26; Available from: http://arxiv.org/abs/1810.11212

24. Hämäläinen MS, Ilmoniemi RJ. Interpreting magnetic fields of the brain: minimum norm estimates. Medical & Biological Engineering & Computing. 1994;32(1):35–42.

25. Pascual-marqui RD. Review of Methods for Solving the EEG Inverse Problem. 1999;1(1):75–86.

26. Srinivasan R. Methods to Improve the Spatial Resolution of EEG [Internet]. Vol. 1, INTERNATIONAL JOURNAL OF BIOELECTROMAGNETISM. 1999. Available from: www.tut.fi/ijbem/

27. Babiloni F, Cincotti F, Carducci F, Rossini PM, Babiloni C. Spatial enhancement of EEG data by surface Laplacian estimation: the use of magnetic resonance imaging-based head models. Clinical Neurophysiology [Internet]. 2001 May;112(5):724–7. Available from: https://linkinghub.elsevier.com/retrieve/pii/S1388245701004941

28. He B, Astolfi L, Valdes-Sosa PA, Marinazzo D, Palva SO, Benar C-G, et al. Electrophysiological Brain Connectivity: Theory and Implementation. IEEE Transactions on Biomedical Engineering [Internet]. 2019 Jul 1;66(7):2115–37. Available from: https://ieeexplore.ieee.org/document/8708690/

29. WHO. World health organization report about preterm birth [Internet]. 2018 [cited 2021 Oct 17]. Available from: http://www.who.int/en/news-room/fact-sheets/detail/preterm-birth

30. Gao W, Lin W, Grewen K, Gilmore JH. Functional Connectivity of the Infant Human Brain. The Neuroscientist [Internet]. 2017 Apr 7;23(2):169–84. Available from: http://journals.sagepub.com/doi/10.1177/1073858416635986

31. Volpe JJ. Brain injury in premature infants: a complex amalgam of destructive and developmental disturbances. The Lancet Neurology [Internet]. 2009 Jan;8(1):110–24. Available from: https://linkinghub.elsevier.com/retrieve/pii/S1474442208702941

32. Himpens E, van den Broeck C, Oostra A, Calders P, Vanhaesebrouck P. Prevalence, type, distribution, and severity of cerebral palsy in relation to gestational age: a meta-analytic review. Developmental Medicine & Child Neurology [Internet]. 2008 May;50(5):334–40. Available from: https://onlinelibrary.wiley.com/doi/10.1111/j.1469-8749.2008.02047.x

33. Spittle AJ, Morgan C, Olsen JE, Novak I, Cheong JLY. Early Diagnosis and Treatment of Cerebral Palsy in Children with a History of Preterm Birth. Clinics in Perinatology [Internet]. 2018 Sep 1;45(3):409–20. Available from: https://linkinghub.elsevier.com/retrieve/pii/S0095510818313708

34. Stoelhorst GMSJ, Rijken M, Martens SE, van Zwieten PHT, Feenstra J, Zwinderman AH, et al. Developmental outcome at 18 and 24 months of age in very preterm children: a cohort study from 1996 to 1997. Early Human Development [Internet]. 2003 Jun;72(2):83–95. Available from: https://linkinghub.elsevier.com/retrieve/pii/S0378378203000112

35. van Beek PE, van der Horst IE, Wetzer J, van Baar AL, Vugs B, Andriessen P. Developmental Trajectories in Very Preterm Born Children Up to 8 Years: A Longitudinal Cohort Study. Frontiers in Pediatrics [Internet]. 2021 May 10;9. Available from: https://www.frontiersin.org/articles/10.3389/fped.2021.672214/full

36. Woodward LJ, Edgin JO, Thompson D, Inder TE. Object working memory deficits predicted by early brain injury and development in the preterm infant. Brain [Internet]. 2005 Nov 1;128(11):2578–87. Available from: http://academic.oup.com/brain/article/128/11/2578/339575/Object-working-memory-deficits-predicted-by-early

37. Leversen KT, Sommerfelt K, Rønnestad A, Kaaresen PI, Farstad T, Skranes J, et al. Predicting neurosensory disabilities at two years of age in a national cohort of extremely premature infants. Early Human Development [Internet]. 2010 12 Sep;86(9):581–6. Available from: https://linkinghub.elsevier.com/retrieve/pii/S0378378210001829

38. Lee ES, Yeatman JD, Luna B, Feldman HM. Specific language and reading skills in school-aged children and adolescents are associated with prematurity after controlling for IQ. Neuropsychologia [Internet]. 2011 Apr;49(5):906–13. Available from: https://linkinghub.elsevier.com/retrieve/pii/S0028393210005828

39. Aylward GP. Neurodevelopmental Outcomes of Infants Born Prematurely. Journal of Developmental & Behavioral Pediatrics [Internet]. 2014 Jul;35(6):394–407. Available from: https://journals.lww.com/00004703-201407000-00007

40. Mangin KS, Horwood LJ, Woodward LJ. Cognitive Development Trajectories of Very Preterm and Typically Developing Children. Child Development [Internet]. 2017 Jan 1;88(1):282–98. Available from: https://onlinelibrary.wiley.com/doi/10.1111/cdev.12585

41. Carmo ALS do, Fredo FW, Bruck I, Lima J do RM de, Janke RNRGH, Fogaça T da GM, et al. Neurological, cognitive and learning evaluation of students who were born preterm. Revista Paulista de Pediatria [Internet]. 2022;40:e2020252. Available from: http://www.scielo.br/scielo.php?script=sci_arttext&pid=S0103-05822022000100406&tlng=en

42. Nunez PL, Silberstein RB, Cadusch PJ, Wijesinghe RS, Westdorp AF, Srinivasan R. A theoretical and experimental study of high resolution EEG based on surface Laplacians and cortical imaging. Electroencephalography and Clinical Neurophysiology [Internet]. 1994 Jan;90(1):40–57. Available from: https://linkinghub.elsevier.com/retrieve/pii/0013469494901120

43. Burle B, Spieser L, Roger C, Casini L, Hasbroucq T, Vidal F. Spatial and temporal resolutions of EEG: Is it really black and whiteã A scalp current density view. International Journal of Psychophysiology. 2015 Sep 1;97(3):210–20.

44. Colclough GL, Brookes MJ, Smith SM, Woolrich MW. NeuroImage A symmetric multivariate leakage correction for MEG connectomes. NeuroImage [Internet]. 2015;117:439–48. Available from: http://dx.doi.org/10.1016/j.neuroimage.2015.03.071

45. Gonzalez-Moreira E, Paz-Linares D, Areces-Gonzalez A, Wang Y, Li M, Harmony T, et al. Bottom-up control of leakage in spectral electrophysiological source imaging via structured sparse bayesian learning. bioRxiv 2020.02.25.964684; Available from: https://doi.org/10.1101/2020.02.25.964684

46. Paz-Linares D, Gonzalez-Moreira E, Martinez-Montes E, Valdes-Hernandez PA, Bosch-Bayard J, Bringas-Vega ML, et al. Caulking the “leakage effect” in MEEG source connectivity analysis. arXiv. 2018.

47. Paz-Linares D, Vega-Hernández M, Rojas-López PA, Valdés-Hernández PA, Martínez-Montes E, Valdés-Sosa PA. Spatio Temporal EEG Source Imaging with the Hierarchical Bayesian Elastic Net and Elitist Lasso Models. Frontiers in Neuroscience [Internet]. 2017 Nov 16;11(NOV). Available from: http://journal.frontiersin.org/article/10.3389/fnins.2017.00635/full13

48. Wipf D, Nagarajan S. A unified Bayesian framework for MEG/EEG source imaging. NeuroImage [Internet]. 2009 Feb 1;44(3):947–66. Available from: https://linkinghub.elsevier.com/retrieve/pii/S1053811908001870

49. Oostenveld R, Praamstra P. The five percent electrode system for high-resolution EEG and ERP measurements. Clinical Neurophysiology [Internet]. 2001 Apr;112(4):713–9. Available from: https://linkinghub.elsevier.com/retrieve/pii/S1388245700005277

50. Fuchs M, Kastner J, Wagner M, Hawes S, Ebersole JS. A standardized boundary element method volume conductor model. Clinical Neurophysiology [Internet]. 2002 May;113(5):702–12. Available from: https://linkinghub.elsevier.com/retrieve/pii/S1388245702000305

51. Tzourio-Mazoyer N, Landeau B, Papathanassiou D, Crivello F, Etard O, Delcroix N, et al. Automated Anatomical Labeling of Activations in SPM Using a Macroscopic Anatomical Parcellation of the MNI MRI Single-Subject Brain. NeuroImage [Internet]. 2002 Jan;15(1):273–89. Available from: https://linkinghub.elsevier.com/retrieve/pii/S1053811901909784

52. Stam CJ, Nolte G, Daffertshofer A. Phase lag index: Assessment of functional connectivity from multi channel EEG and MEG with diminished bias from common sources. Human Brain Mapping [Internet]. 2007 Nov;28(11):1178–93. Available from: https://onlinelibrary.wiley.com/doi/10.1002/hbm.20346

53. Vinck M, Oostenveld R, van Wingerden M, Battaglia F, Pennartz CMA. An improved index of phase-synchronization for electrophysiological data in the presence of volume-conduction, noise and sample-size bias. NeuroImage. 2011 Apr 15;55(4):1548–65.

54. Cohen MX. Analyzing neural time series data: Theory and practice. 2014.

55. Achard S, Bullmore E. Efficiency and Cost of Economical Brain Functional Networks. Friston KJ, editor. PLoS Computational Biology [Internet]. 2007 Feb 2;3(2):e17. Available from: https://dx.plos.org/10.1371/journal.pcbi.0030017

56. Wozniak JR, Mueller BA, Bell CJ, Muetzel RL, Hoecker HL, Boys CJ, et al. Global functional connectivity abnormalities in children with fetal alcohol spectrum disorders. Alcoholism, clinical and experimental research [Internet]. 2013 May;37(5):748–56. Available from: http://www.ncbi.nlm.nih.gov/pubmed/23240997

57. Cleveland WS, Devlin SJ. Locally Weighted Regression: An Approach to Regression Analysis by Local Fitting Locally Weighted Regression: An Approach to Regression Analysis by Local Fifing. Vol. 83, Source: Journal of the American Statistical Association. 1988.

58. Dereymaeker A, Pillay K, Vervisch J, de Vos M, van Huffel S, Jansen K, et al. Review of sleep-EEG in preterm and term neonates. Early Human Development [Internet]. 2017 Oct 1;113:87–103. Available from: https://linkinghub.elsevier.com/retrieve/pii/S037837821730325014

59. Gao W, Alcauter S, Elton A, Hernandez-Castillo CR, Smith JK, Ramirez J, et al. Functional network development during the first year: Relative sequence and socioeconomic correlations. Cerebral Cortex. 2015 Sep 1;25(9):2919–28.

60. Gao W, Alcauter S, Smith JK, Gilmore JH, Lin W. Development of human brain cortical network architecture during infancy. Brain Structure and Function. 2015 Mar 1;220(2):1173–86.

61. Toulmin H, Beckmann CF, O’Muircheartaigh J, Ball G, Nongena P, Makropoulos A, et al. Specialization and integration of functional thalamocortical connectivity in the human infant. Proceedings of the National Academy of Sciences of the United States of America [Internet]. 2015 May 19;112(20):6485–90. Available from: http://www.ncbi.nlm.nih.gov/pubmed/25941391

62. Smyser CD, Inder TE, Shimony JS, Hill JE, Degnan AJ, Snyder AZ, et al. Longitudinal analysis of neural network development in preterm infants. Cerebral cortex (New York, NY : 1991) [Internet]. 2010 Dec;20(12):2852–62. Available from: http://www.ncbi.nlm.nih.gov/pubmed/20237243

63. Thompson DK, Chen J, Beare R, Adamson CL, Ellis R, Ahmadzai ZM, et al. Structural connectivity relates to perinatal factors and functional impairment at 7 years in children born very preterm. NeuroImage [Internet]. 2016 Jul 1;134:328–37. Available from: https://linkinghub.elsevier.com/retrieve/pii/S1053811916300143

64. Rogers CE, Lean RE, Wheelock MD, Smyser CD. Aberrant structural and functional connectivity and neurodevelopmental impairment in preterm children. Vol. 10, Journal of Neurodevelopmental Disorders. BioMed Central Ltd.; 2018.

65. Thatcher RW, Biver CJ, North D. Spatial-Temporal Current Source Correlations and Cortical Connectivity. Clinical EEG and Neuroscience [Internet]. 2007 Jan 25;38(1):35–48. Available from: http://journals.sagepub.com/doi/10.1177/155005940703800109

66. Meijer EJ, Hermans KHM, Zwanenburg A, Jennekens W, Niemarkt HJ, Cluitmans PJM, et al. Functional connectivity in preterm infants derived from EEG coherence analysis. European Journal of Paediatric Neurology. 2014 Nov 1;18(6):780–9.

67. González JJ, Mañas S, de Vera L, Méndez LD, López S, Garrido JM, et al. Assessment of electroencephalographic functional connectivity in term and preterm neonates. Clinical Neurophysiology. 2011 Apr;122(4):696–702.

68. Hinojosa-Rodríguez M, Harmony T, Carrillo-Prado C, van Horn JD, Irimia A, Torgerson C, et al. Clinical neuroimaging in the preterm infant: Diagnosis and prognosis. NeuroImage: Clinical [Internet]. 2017;16:355–68. Available from: https://linkinghub.elsevier.com/retrieve/pii/S2213158217302061

69. Harmony T. Early diagnosis and treatment of infants with prenatal and perinatal risk factors for brain damage at the neurodevelopmental research unit in Mexico. 15 NeuroImage [Internet]. 2021 Jul 15;235:117984. Available from: https://linkinghub.elsevier.com/retrieve/pii/S1053811921002615

70. Sakai T, Mikami A, Suzuki J, Miyabe-Nishiwaki T, Matsui M, Tomonaga M, et al. Developmental trajectory of the corpus callosum from infancy to the juvenile stage: Comparative MRI between chimpanzees and humans. PLoS ONE. 2017 Jun 1;12(6).

71. Keshavan MS, Diwadkar VA, DeBellis M, Dick E, Kotwal R, Rosenberg DR, et al. Development of the corpus callosum in childhood, adolescence and early adulthood. Life Sciences [Internet]. 2002 Mar;70(16):1909–22. Available from: https://linkinghub.elsevier.com/retrieve/pii/S0024320502014923

72. Garvey MA, Mall V. Transcranial magnetic stimulation in children. Clinical Neurophysiology [Internet]. 2008 May;119(5):973–84. Available from: https://linkinghub.elsevier.com/retrieve/pii/S1388245707007353

73. Ciechanski P, Zewdie E, Kirton A. Developmental profile of motor cortex transcallosal inhibition in children and adolescents. Journal of Neurophysiology [Internet]. 2017 Jul 1;118(1):140–8. Available from: https://www.physiology.org/doi/10.1152/jn.00076.2017

74. Hasegawa T, Yamada K, Morimoto M, Morioka S, Tozawa T, Isoda K, et al. Development of Corpus Callosum in Preterm Infants Is Affected by the Prematurity: In Vivo Assessment of Diffusion Tensor Imaging at Term-Equivalent Age. Pediatric Research [Internet]. 2011 Mar;69(3):249–54. Available from: http://www.nature.com/doifinder/10.1203/PDR.0b013e3182084e54

75. Huttenlocher PR, Dabholkar AS. Regional differences in synaptogenesis in human cerebral cortex. The Journal of Comparative Neurology [Internet]. 1997 Oct 20;387(2):167–78. Available from: https://onlinelibrary.wiley.com/doi/10.1002/(SICI)1096-9861(19971020)387:2<167::AID-CNE1>3.0.CO;2-Z

76. Lebenberg J, Mangin JF, Thirion B, Poupon C, Hertz-Pannier L, Leroy F, et al. Mapping the asynchrony of cortical maturation in the infant brain: A MRI multi-parametric clustering approach. NeuroImage. 2019 Jan 15;185:641–53.

77. Paus T, Collins DL, Evans AC, Leonard G, Pike B, Zijdenbos A. Maturation of white matter in the human brain: a review of magnetic resonance studies. Brain Research Bulletin [Internet]. 2001 Feb;54(3):255–66. Available from: https://linkinghub.elsevier.com/retrieve/pii/S0361923000004342

78. Woodward LJ, Clark CAC, Pritchard VE, Anderson PJ, Inder TE. Neonatal White Matter Abnormalities Predict Global Executive Function Impairment in Children Born Very Preterm. Developmental Neuropsychology [Internet]. 2011 Jan 25;36(1):22–41. Available from: http://www.tandfonline.com/doi/abs/10.1080/87565641.2011.540530

79. Otero GA, Harmony T, Pliego-Rivero FB, Ricardo-Garcell J, Bosch-Bayard J, Porcayo-Mercado R, et al. QEEG norms for the first year of life. Early Human Development [Internet]. 2011 Oct;87(10):691–703. Available from: https://linkinghub.elsevier.com/retrieve/pii/S037837821100200316

80. Roca-Stappung M, Moguel-González M, Fernández T, Harmony T, Mendoza-Montoya O, Marroquín JL, et al. Characterization of the Sensorimotor Rhythm in 4-Month-Old Infants Born at Term and Premature. Applied Psychophysiology Biofeedback. 2017 Dec 1;42(4):257–67.

